# Vitrocam: A simple low cost Vitrobot camera for assessing grid quality

**DOI:** 10.1101/2022.06.16.496351

**Authors:** Eugene Y.D. Chua, Viacheslav Serbynovskyi, Robert Gheorghita, Lambertus M. Alink, Daniel Podolsky, Clinton S. Potter, Bridget Carragher

## Abstract

The most widely used sample preparation method for single particle cryo-electron microscopy (cryo-EM) today involves the application of 3-4 μl of sample onto a cryo-EM grid, removing most of the liquid by blotting with filter paper, then rapidly plunging into liquid ethane to vitrify the sample. To determine if the grid has appropriate ice thicknesses and sufficient area for cryo-EM imaging, the grid must be inserted into a transmission electron microscope (TEM) and screened. This process to evaluate grid quality is costly and time consuming. Here, we present our initial attempt to image the sample preparation process in one of the most commonly used plunge freezing devices, the Vitrobot. We do this by building the Vitrocam, a Raspberry Pi high-speed camera, that captures images of grids mid-plunge. Images from the Vitrocam can be correlated to TEM atlases and show promise for providing preliminary feedback on grid quality and ice thickness.

## Introduction

Plunge freezing as a method for sample preparation for cryo-electron microscopy (cryo-EM) was first developed in the 1980s ^1–3^. This method involves applying a 3-4 μl volume of sample to a hydrophilized grid, removing most of the liquid by blotting with filter paper to leave behind a very thin film of liquid, then rapidly plunging the grid into a cryogen, typically liquid ethane. The vitrification of the liquid preserves the biological sample in a state that is suitable for cryo-EM imaging.

Today, there are commercial devices available that perform this task, including the Vitrobot (ThermoFisher Scientific), Leica’s EM GP (Leica), and Gatan’s CP3 (Gatan); the Vitrobot is the most widely used of these instruments. These devices are generally equipped with a humidity- and temperature-controlled blotting chamber, mechanical blotting pads that can be adjusted for blot force and time, and in some cases temperature control for liquid ethane. However, this method of plunge freezing poses several challenges. Firstly, controlling ice thickness is difficult. These devices use large-scale mechanical forces in an attempt to create thin films of liquid on the order of 100 nm ^4^. As a result, there can be wide variations in the resulting ice thicknesses across a single grid, and it can be difficult to exactly reproduce the same ice thickness between grids. Secondly, the lack of any feedback during the plunge freezing process means that in order to evaluate the quality of ice on a grid, the grid must be clipped (if using an autoloader system), loaded onto a transmission electron microscope (TEM), an atlas collected, and several higher magnification exposure images taken to assess ice thickness and sample quality. The cost of a grid cartridge for an autoloader system is ^~^$20 per grid, and screening microscope time costs $50-100 an hour, with each grid requiring ^~^30-45 minutes to load, acquire an atlas, and screen. Furthermore, staff time is required at each of these steps. Occasionally, grids loaded onto a TEM are discovered to be mostly electron opaque because the ice is too thick, or they may be bent out of shape or contain many broken squares from poor handling. The scientist must then switch to the next grid in hopes of finding a suitable one, or return to the plunge freezer for another round of grid preparation and screening. This iterative process could be improved, and costs reduced, by providing more feedback about grid quality during the sample preparation stage, rather than waiting until TEM imaging.

There exist several grid preparation devices that have implemented grid quality assessment feedback during the preparation stage. The Spotiton ^5,6^ and chameleon ^7,8^ vitrification devices use piezoelectric dispensers to apply sample onto self-wicking grids, which wick away excess solution to leave behind a thin film of liquid on the grid ^9,10^. Spotiton and chameleon incorporate high-speed cameras that visualize the wicking process and provide feedback for determining the grid’s suitability for transferring to the TEM for imaging. The camera acquires images of the grid during the wicking process, and by examining the change in square sizes during wicking, an experienced user can estimate the likely ice thickness outcome in a TEM. At the very least, it is possible to tell when a grid did not successfully wick and that it will have ice that is too thick and thus should be discarded. Recently, a novel sample preparation device coupled to a Linkam stage ^11^ was developed that utilizes a light microscope to monitor the liquid aspiration process and determine the right moment for vitrification, so to improve control and reproducibility of the resulting ice thicknesses. All these devices have thus incorporated cameras into the grid preparation process to improve success rates and limit wasting time and money on grids that are not suitable for imaging.

Here, we describe a simple, low-cost camera system we call the Vitrocam to assess grid quality during plunge freezing in a Vitrobot. Using a Raspberry Pi coupled to a high-speed camera, we capture grid images mid-plunge, after being blotted and shortly before vitrification in liquid ethane, to obtain preliminary information about grid quality and liquid thickness. While there are still several improvements that need to be made to the system, the Vitrocam shows promise as a device that can provide useful information on grid quality at the sample preparation stage, thereby saving time, resources, and money. We are putting this early description of the Vitrocam onto bioRxiv so that other groups will be encouraged to try it out and develop it further.

## Materials and methods

### Equipment setup

The centerpiece of the Vitrocam is a high-speed OV9281 camera that is controlled by a Raspberry Pi 4 Model B. We have also added an infra-red (IR) proximity sensor to detect the position of the blotting pads, and a pair of IR LED lights for illumination (see Supplementary Table 1 for a parts list). The Pi is held outside the Vitrobot, and the jumper cables connecting the Pi to the camera, IR proximity sensor, and IR LEDs are fed through the left sample port on the Vitrobot chamber (Figure 1). In addition, an external light source (an Amscope 150W Fiber Optic Dual Gooseneck Microscope Illuminator, in this case) is used to provide illumination to the camera (Figure 1).

**Figure 1.**
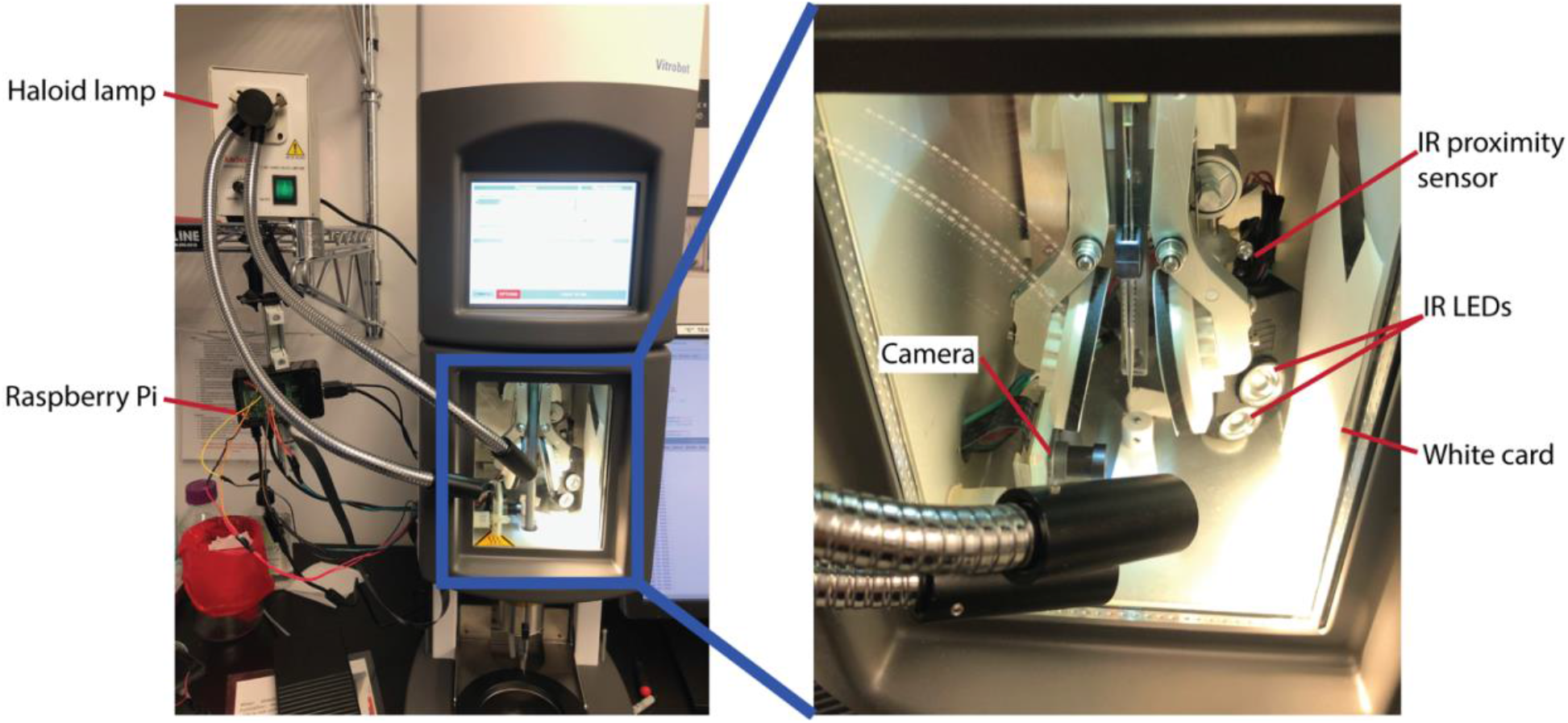
Vitrocam setup, consisting of a Raspberry Pi located outside the Vitrobot, connected via the left sample port to a high-speed OV9281 camera, an IR proximity sensor, and a pair of IR LED lights, all mounted within the Vitrobot chamber. A haloid lamp (Amscope 150W Fiber Optic Dual Gooseneck Microscope Illuminator) provides additional lighting, and a white card covering the right sample port provides even illumination across the camera field of view.

The OV9281 camera sensor was selected for several important features: it can capture images at >200 frames per second, has a global (as opposed to rolling) shutter, has a black and white sensor, and can be controlled by the Rasberry Pi. The camera came with a wide-angle fisheye lens (Fov(D)=148 degrees, Fov(H)=118 degrees, Focal length ^~^2.8 mm) on an M12 mount. We tested a variety of lenses, but the stock lens was the most suitable. The camera is held in place in the Vitrobot chamber with a custom 3D-printed holder that is attached to the wall of the chamber with Velcro. The important feature of the holder is that it holds the camera firmly in place. Due to the narrow depth of focus of the camera, any change in position of the camera, e.g. by opening and closing the doors to the chamber, will result in out-of-focus images. The camera was positioned and focused on a grid attached to the Vitrobot tweezers in the ‘Process’ position. Placing the camera right on the edge of the tweezer opening allows for the closest possible view of the grid. This must be done carefully to prevent any potential collisions with and damage to the tweezers and Vitrobot.

The IR proximity sensor is positioned behind the arm of the right blotting pad with the help of a 3D-printed holder. The sensitivity and position of the proximity sensor are adjusted such that it will detect the presence or absence of the blotting pad arm during resting or blotting respectively. These adjustments can be made by using the indicator lights on the proximity sensor, or with the help of a simple script on the Pi reporting the detection of the blotting pad arm.

A matte white card large enough to cover the field of view of the camera (^~^7 cm x 10 cm) is placed on the right wall of the Vitrobot chamber. The white card serves to cover up the sample port and provides a surface for creating even illumination. An intense light source is needed to give sufficient illumination to the camera. Here, we used an Amscope 150W Fiber Optic Dual Gooseneck Microscope Illuminator that was available in-house. The lights point directly at the white card from outside the Vitrobot chamber through the glass door. The positions of the lights are adjusted to give as even illumination as possible, as seen through the camera. In addition, IR LEDs for providing IR illumination to the camera are positioned below the IR proximity sensor on the same holder. The LEDs point towards the card at an angle and must be positioned to give maximum even illumination to the camera. Sufficient and even illumination for both the IR LEDs and external light source are determined by using the camera at the capture settings of ^~^200 fps and 30 μs shutter speed.

### Software setup

Our Raspberry Pi 4 is running Raspbian GNU/Linux 10 (Buster). The OV9281 camera is controlled by the Pi’s native libcamera software suite. Automatic video capture of the plunging process is achieved by using a simple Python script that connects the status of the IR proximity detectors to the video capture (Python script is available on https://github.com/eydchua/vitrocam). Firstly, the script monitors the status of the blotting pad arms through the IR proximity sensor. When blotting begins, the blotting pad arms move out of range of the IR proximity sensor, triggering a “Blotting in progress” report in the script. When blotting is complete, the blotting pad arms move back to the resting position and are detected once more by the IR proximity sensor. This triggers the capture of a short video of the plunge, which is stored in a date-time-stamped folder, and the frames extracted into the same folder. The script then proceeds to a waiting loop where it continuously checks for the next blotting cycle. In this way, the script only needs to be started at the beginning of a Vitrobot session and will automatically capture all the grids prepared into the unique date-time-stamped folders.

## Results and Discussion

### Images of grids can be captured mid-plunge and correlated with TEM atlases

Using Quantifoil R1.2/1.3 300 or 400 mesh grids, we froze water or apoferritin in the Vitrobot with the Vitrocam in operation. At ^~^200 fps and with only the Haloid lamp for illumination, the Vitrocam successfully captured 1-2 frames in which the grid was within the field of view (Figure 2a). Several features were easy to visualize with this Vitrocam setup: (1) when the ice is sufficiently thick, and there is a gradient, it is readily visible (Figure 2b, top two rows); (2) broken squares are readily visible (Figure 2b, bottom row); (3) due to the narrow focal depth of the camera, a grid that is significantly bent will become out of focus. Imaging the grids that we prepared in a TEM, we found that, when there are distinguishing features on the grid such as a large gradient of ice thickness or broken squares, we could readily correlate the images from the Vitrocam with the TEM atlases (Figure 2).

**Figure 2.**
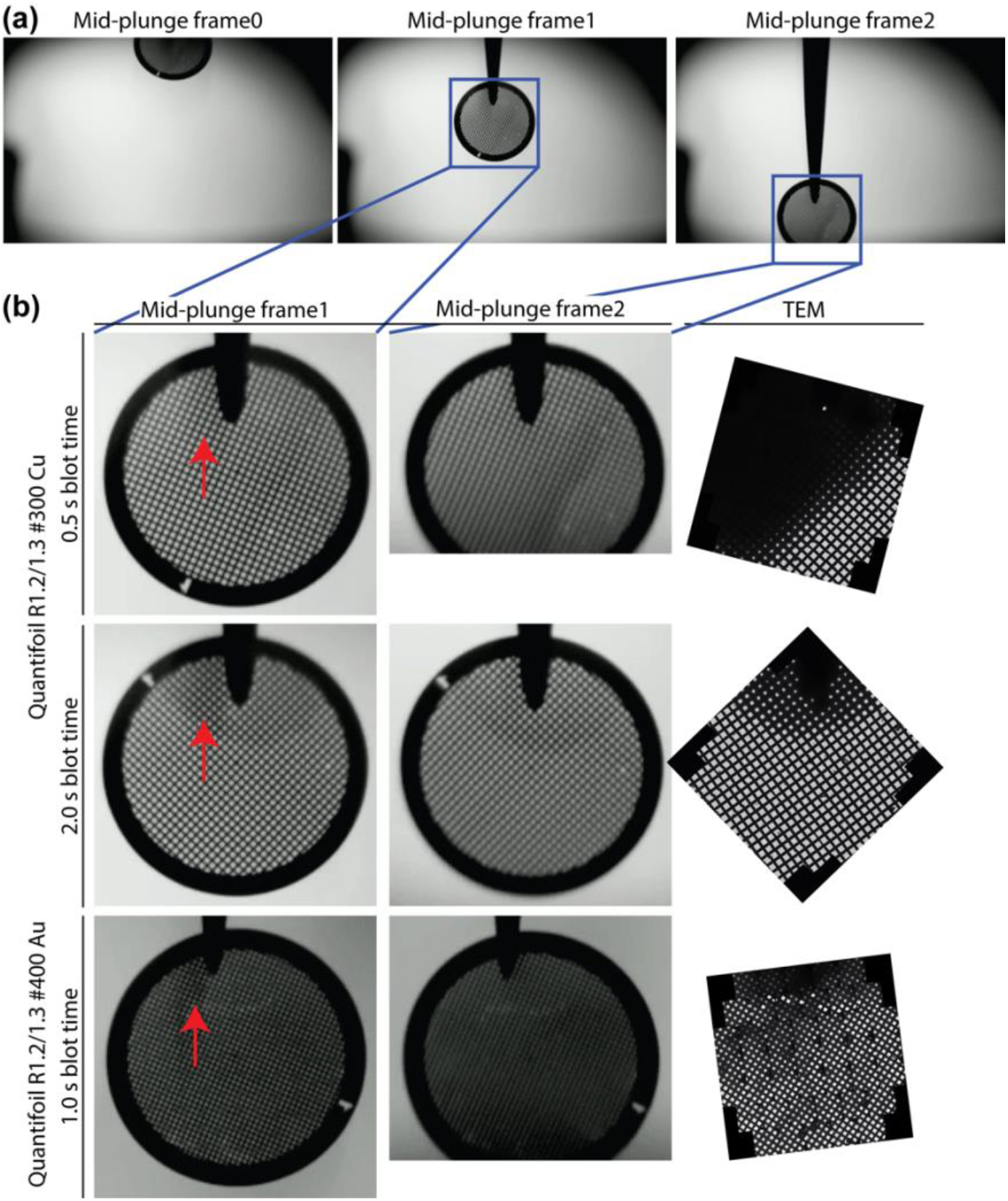
Images of grids captured mid-plunge, and their corresponding TEM atlases. (a) Three consecutive raw frames extracted from a Vitrocam video capturing a grid during the plunging process. (b) Quantifoil grids with 300 (top two rows) or 400 (bottom row) mesh sizes were used, and Vitrocam frames containing the grid are shown along with the corresponding atlas obtained in the TEM. Some images were truncated because the grid had moved out of the field of view. Red arrows indicate a shadow, probably from the tweezers, cast by the lighting setup.

### Challenges with illumination

A fast shutter speed on the camera was needed to capture still images of the grids mid-plunge; however, this then required the use of a sufficiently bright light source for illumination. During our development of the Vitrocam, we used an in-house 150 W Haloid lamp for illumination as a temporary solution. While this setup provided sufficient brightness, the illumination was not even across the field of view (Figure 2a), which resulted in differences in the amount of contrast or information depending on the position of the grid within the camera’s field of view. We also noticed that a shadow, probably from the tweezers, was sometimes cast onto the grid (Figure 2b, red arrows). This was not from the existing LED light in the Vitrobot chamber, but instead probably originated from our new lighting setup. We envision that having a light source directly facing the camera, as in the manner of Spotiton and chameleon’s grid cameras, will result in fewer shadows and more even illuination.

The uneven illumination from the Haloid lamp means that the same grid seen in two different frames can look different (Figure 2, compare ‘Mid-plunge frame1’ with ‘Mid-plunge frame2’). Having better and more even illumination across the camera’s field of view will be important for more objective assessments of liquid thicknesses on grids. Once enough Vitrocam data is collected and coupled to TEM ice thickness information, we will be able to provide more accurate estimations of the expected ice thicknesses based on the Vitrocam images.

Currently, obtaining even illumination across the field of view is complicated by (1) the large field of view of the camera, due to the fisheye lens; (2) limited space in the Vitrobot chamber; and (3) requirement for access via the right sample port to apply sample to the grid in the ‘Process’ position. We are currently testing and improving this setup to overcome these challenges, including changing from a fisheye to a magnifying lens, and assembling an LED setup that is small enough to fit into the Vitrobot chamber but also large enough to illuminate the entire field of view evenly. We are also considering different options for allowing sample application via the right sample port, while not compromising on image quality on the Vitrocam.

### Using IR light enables better visualization of liquid thicknesses

The absorption of light by water in the IR range is 10^2^ to 10^6^-fold better than in the visible light range (Figure 3a). We thus reasoned that having IR illumination will provide more information about liquid thickness on the grid. Preliminary results show that using the IR LEDs in our current Vitrocam setup provides better information about liquid thickness when 3 μl of sample is applied to the grid, compared to just using the Haloid lamp (Figure 3b, ‘After sample application’, comparing ‘Haloid lamp only’ with ‘IR LED only’). Liquid thickness gradients can also be seen with IR LEDs alone (Figure 3b, bottom row), which is a promising outcome for future developments. The amount of additional information about liquid thicknesses that can be gained from using the IR LEDs as opposed to the Haloid lamp on the Vitrocam remains to be determined.

**Figure 3.**
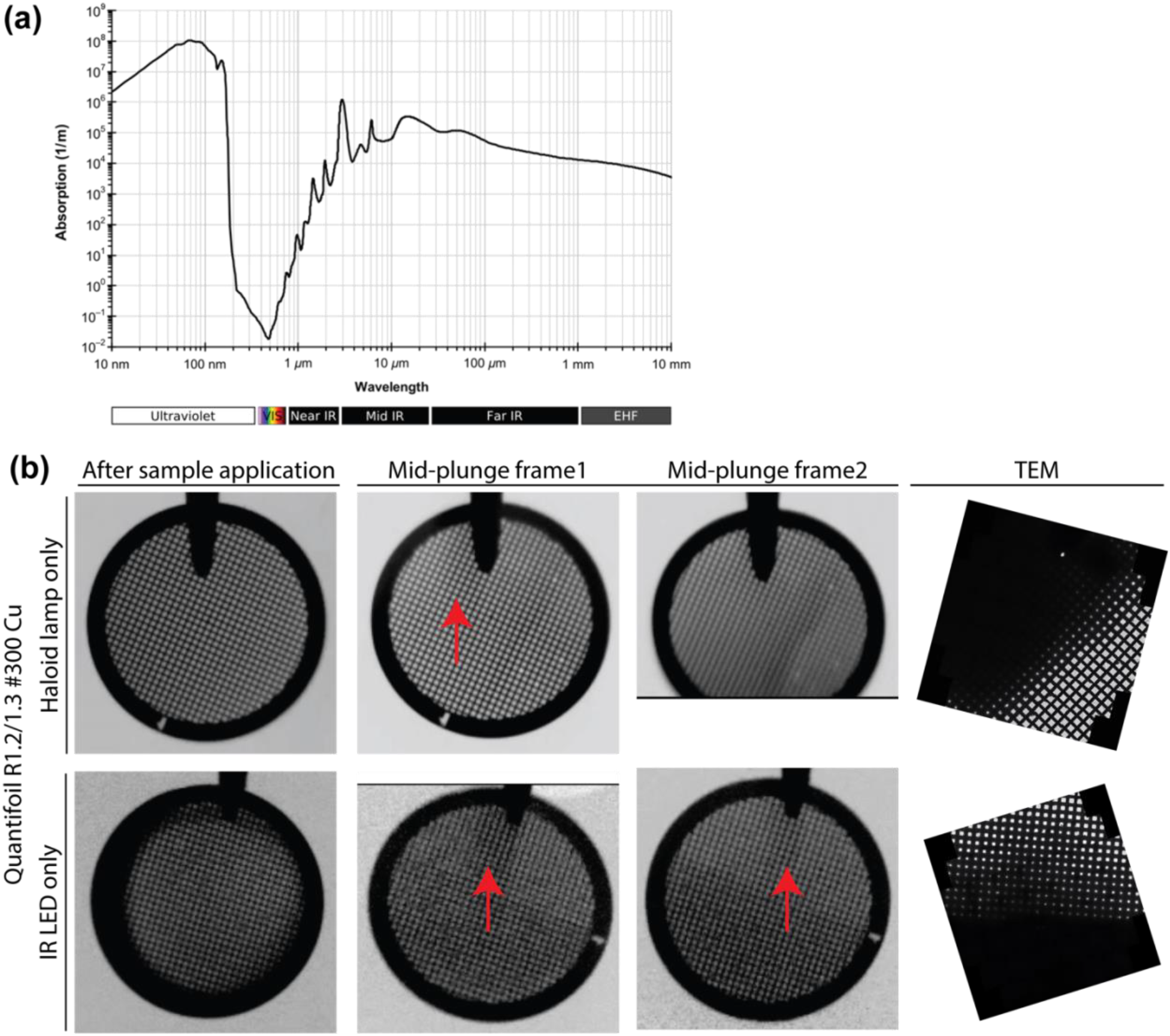
Visualization of grids using white and infra-red illumination. (a) Absorption spectrum of water. Image from Kebes at English Wikipedia, CC BY-SA 3.0, https://commons.wikimedia.org/w/index.php?curid=23793083. (b) Comparison of Haloid lamp illumination (top row) and IR LED illumination (bottom row). Red arrows point to shadows, probably from the tweezers, cast by the lighting setup.

One point of note for the IR setup is that to image gold foil grids, the intensity of the light source must be increased compared to the intensity required to image carbon foil grids (Figure 4).

**Figure 4.**
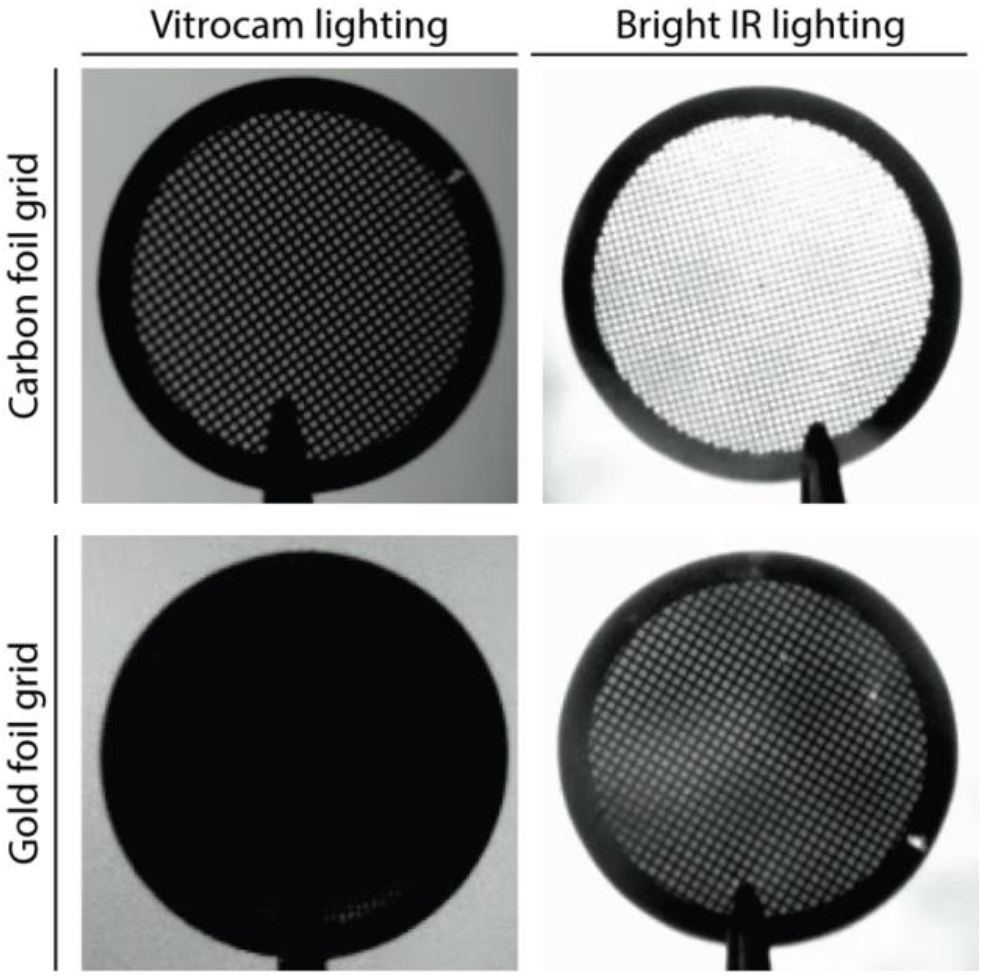
Gold foil grids under initial Vitrocam lighting conditions appear black (bottom left quadrant). Increasing the brightness of an IR LED source placed behind the grid directly facing the camera results in visible squares (bottom right quadrant).

### Vitrocam costs are minimal

The current setup for the Vitrocam (not including the Haloid lamp, monitor, keyboard, and mouse), costs ^~^$215 (see Supplementary Table 1). A new Amscope Haloid lamp is available for ^~^$250, but we envision that this will be superseded by a Pi-controlled lighting setup in future versions of the Vitrocam. A computer monitor, mouse, and keyboard are required for controlling the Pi, and considering their general availability, have not been included in the cost calculations. One can also set up the Pi with VPN access, which will allow the use of any existing computer to control the Vitrocam.

### Improving video capture automation

The current automation setup relies heavily on the proximity sensor to detect the position of the blotting pad arms. While this setup is reliable, it is unable to obtain frame-perfect timing for capturing the grid mid-plunge. For this reason, we capture a short 0.5 second video then analyze the output frame-by-frame. Obtaining frame-perfect timing will be possible by coupling video capture with the Vitrobot software. Alternatively, a simple computer vision program should be able to automatically select frames from our short 0.5 second video that contain the grid.

## Summary

The Vitrocam is a compact and economic setup to capture images of grids during plunge freezing to provide feedback on grid quality, and to some extent, ice thickness. Several improvements to the system are in progress: most crucially we are developing a dedicated IR LED setup as a light source, and in addition will add a suitable magnifying lens for the camera. The cost-effectiveness of the Vitrocam, its ease of assembly, and the ability to install it without making any hardware modifications to the Vitrobot (thereby avoiding any warranty issues), makes it a useful tool for any laboratory with a Vitrobot. By effectively utilizing the ‘sneak-preview’ information gained via the Vitrocam, a significant amount of time and money may be saved by throwing out grids that are obviously bad prior to clipping and imaging.

## Acknowledgements

We thank Kasahun Neselu and Jing Wang at the Simons Electron Microscopy Center for being alpha testers of the Vitrocam. This work was supported by the Simons Electron Microscopy Center and National Resource for Automated Molecular Microscopy located at the New York Structural Biology Center, supported by grants from the Simons Foundation (SF349247) and the NIH National Institute of General Medical Sciences (GM103310).

---

**Supplementary Table 1.**

| Item | Supplier/Manufacturer | Cost (USD) |
| --- | --- | --- |
| CanaKit Raspberry Pi 4 Extreme Kit | CanaKit | 139.99 |
| OV9281 black and white CMOS sensor | Innomaker | 59.99 |
| IR proximity sensor | OSOYOO | 0.99 |
| IR LEDs | DORHEA | 6.99 |
| Jumper wires | EDGELEC | 1.00 |
| 24" camera cable | Adafruit | 6.41 |
|  | Total cost | 215.37 |

## Additional items

- [Amscope 150W Fiber Optic Dual Gooseneck Microscope Illuminator](https://amscope.com/products/hl250-ay?gclsrc=aw.ds&gclsrc=ds), $294.99
- Computer monitor, mouse, and keyboard

## Notes

### Competing Interest Statement

The authors have declared no competing interest.

